# Microbial utilisation of maize rhizodeposits applied to an agricultural soil at a range of concentrations

**DOI:** 10.1101/2024.03.15.585161

**Authors:** Daniela Niedeggen, Lioba Rüger, Eva Oburger, Michael Santangeli, Mutez Ahmed, Doris Vetterlein, Sergey Blagodatsky, Michael Bonkowski

**Affiliations:** University of Cologne, Institute for Zoology, Terrestrial Ecology, Cologne, Germany; Cluster of Excellence on Plant Sciences (CEPLAS), University of Cologne, Germany; University of Natural Resources and Life Sciences, Institute of Soil Research, Tulln an der Donau, Austria; Technical University Munich –TUM, School of Life Sciences, Freising, Germany; Helmholtz Centre for Environmental Research GmbH –UFZ, Dept. Soil System Science, Halle/Saale, Germany

**Keywords:** Carbon cycling, Microbial respiration, Microbiome, Mucilage, Rhizodeposition, Rhizosphere, Root Exudates, Soil microbial activity, Zea mays

## Abstract

Rhizodeposition fuels carbon (C) and nutrient cycling in soil. However, the dynamics of microbial growth on rhizodeposits in relation to the distance from the root have not been well studied. This study investigates microbial growth on individual organic components of rhizodeposits and on maize root-derived exudates and mucilage from an agricultural soil. By creating a gradient of substrate concentrations, we simulated reduced microbial access to rhizosphere C with increasing distance to the root surface. We identified distinct C-thresholds for the activation of microbial growth, and these were significantly higher for rhizodeposits compared to singular, simple sugars. In addition, testing for stoichiometric constraints of microbial growth by supplementing N and P showed accelerated and increased microbial growth by activating a larger proportion of the microbial biomass. Early and late season exudates triggered significantly different microbial growth responses. The mineralisation of early season exudates was induced at a high C-threshold, whereas the mineralisation of late season exudates showed ’sugar-like’ properties, with a low C-threshold, high substrate affinity, and a reduced maximum respiration rate. Mucilage exhibited the highest C-threshold for the activation of microbial growth, although with a short lag-period and with an efficient mucilage degradation comparable to that of sugars. By determining kinetic parameters and turnover times for different root-derived substrates, our data enable the upscaling of micro-scale processes to the whole root system, allowing more precise predictions of how rhizodeposits drive microbial C and nutrient dynamics in soil.

**Highlights:**

- Growth thresholds for rhizodeposits were significantly higher than for singular, simple sugars.
- No distinct microbial growth on root exudates was observed even at high concentrations.
- Mucilage has a short lag-phase and efficient decomposition like sugars but only above a high threshold.
- N and P limited microbial growth in the agricultural soil.

## Introduction

More than a century ago, the rhizosphere was defined as the region around roots influenced by rhizodeposition (Hiltner, 1904), but still today the precise boundaries of the rhizosphere remain elusive (Finzi et al., 2015). The release of labile organic C from roots creates temporally a specific hotspot with significantly enhanced microbial activity and growth (Jones et al., 2004; Kuzyakov & Blagodatskaya, 2015; Ma et al., 2018). The microbial activity decreases strongly with distance from the root surface (Vetterlein et al., 2020). The spatial expansion of the rhizosphere is influenced by the amount and composition of rhizodeposits, their diffusion and sorption in soils, which is a function of plant age, photosynthetic activity, root morphology and soil texture, but also depends on their degradation by microbial activity (Hertenberger et al., 2002; Ma et al., 2018; Molina et al., 2001; Santangeli et al., 2024).

Rhizodeposits are typically divided into actively secreted mucilage, lubricating the advancing root tip, and mostly passive released exudates (Nguyen, 2003). Exudates, which are both low-molecular-weight compounds such as sugars, amino acids, organic acids, phenolics, and high molecular weight compounds, like proteins, contribute significantly to the diversity of rhizodeposits (Collins & Reilly, 1968; Lohse et al., 2021; Walter et al., 2003). Less diverse is the mucilage, a viscous substance actively secreted at the root tip, consisting of high molecular weight compounds such as polysaccharides, proteins, phenolic acids and lipids (Bacic, 1987). These rhizodeposits provide the primary energy source for microbial activity in the rhizosphere (Brimecombe, 2000).

While physical factors are integrated into existing rhizosphere models (Kuppe et al., 2022), the consideration of biological factors, such as consumption and mineralization of rhizodeposits by microorganisms remains a challenge. Current models either lack an explicit representation of microbial degradation activities or follow simplified linear degradation kinetics (Kirk et al., 1999; Landl et al., 2021). More advanced models (Chertov et al., 2022; Finzi et al., 2015) need to be rigorously validated against experimental data obtained in the plant-soil environment, as the microbial parameters in rhizosphere models are traditionally derived from pure culture studies (Zelenev et al., 2000). Integrating biological activity into rhizosphere models remains a challenge due to the lack of quantitative data, leaving a gap in the modelling of rhizosphere processes (Schnepf et al., 2022).

Because the vast majority of microorganisms in bulk soil is dormant (Stenström et al., 2001), microbial growth is initiated only 5-15 hours after the C-pulse from rhizodeposition (Anderson & Domsch, 1978; Anderson & Domsch, 1985; Reischke et al., 2015). The transition of microorganisms from a stationary dormant to an active growth stage is associated with a dramatic change in their metabolism. After encountering an accessible C source, the dormant soil microorganisms must first upregulate their enzymatic machinery for mineralising the growth substrates, which is characterized by a typical lag-period with significantly increased microbial respiration before the onset of microbial growth (Panikov, 1995). The lag-period depends on the proportion of dormant microbial biomass, and microbial respiration increases with the concentration of the C source until the “maximum initial respiratory response” (MIRR) is reached at maximum or excess availability of C (Anderson & Domsch, 1978). Microbial mineralization and growth responses have mainly been studied using glucose, a common component of rhizodeposits (Jones et al., 2009). Previous studies have shown that microbial growth in response to glucose is triggered only when the concentration of carbon exceeds a threshold value that depends on the soil microbial biomass (Cmic) (Anderson & Domsch, 1985; Reischke et al., 2015; Sawada et al., 2008). The studies have reported a C-threshold of approximately 50-150% Cmic. However, microbial responses to glucose versus rhizodeposits, the latter being a diverse mixture of different compounds, have not been compared in a single experimental set-up, and microbial growth kinetics on rhizodeposits have not been studied at all. We argue that mineralization dynamics and C-thresholds governing microbial growth in response to rhizodeposition differ from those associated with glucose and require further investigation. In addition, the C-rich rhizodeposits are expected to affect the stoichiometric requirements of the rhizosphere microorganisms, and it is unclear how microbial growth is affected by N- or P-limitation.

This study aims to provide quantitative information on the microbial responses to and mineralisation of natural maize rhizodeposits and of their most common single compounds in an agricultural soil. Microbial responses to single C sources, such as simple sugars, organic acids or amino acids, will be compared with those to complex sources, such as exudates or mucilage, in a single set-up. With high temporal resolution, we measured the kinetics of microbial respiration and growth in response to single C sources, including mono- and polysaccharide sugars, amino and organic acids, as well as complex substrates such as maize root-derived exudates or mucilage, quantifying the microbial growth responses to these different carbon sources. A gradient of substrate concentration was created to simulate reduced microbial access to C with increasing distance from the root surface. Further nutrient limitations were tested by supplementing N and/or P along the concentration gradient at a stoichiometric ratio of C:N:P 10:2:1. For the first time, our study provides information to accurately calibrate models of rhizosphere microbial growth dynamics with increasing distance from the root surface.

## Material and Methods

### Sampling of soil and origin of rhizodeposits

A loamy agricultural soil was sampled in June 2022 from a maize field experiment near Bad Lauchstädt, Germany (see Vetterlein et al. (2021), sieved (2 mm) and stored at 4°C until use. The soil had a pH of 5.6, and a C:N ratio of 10.85, containing 0.80% C and 0.07% N. The extractable organic C (EOC), denoting the potentially available, organic C for microbial growth, was determined by extracting samples of 5 g field-moist soil with 20 ml of 0.025M K_2_SO_4_ on a horizontal shaker at 250 rpm for 30 min. Subsequently, samples were centrifuged for 30 min at 4420 g and EOC was measured using a EOC-TN Analyzer (Multi-N/C 2100S, Analytik Jena, Germany).

The single components of maize rhizodeposits tested in this study, including the sugars glucose (C_6_H_12_O_6_, VWR), sucrose (C_12_H_22_O_11_), and arabinose (C_5_H_10_O_5_), the amino acid aspartate (C_4_H_7_NO_4_, Serva), and the organic acids citric acid (C_6_H_8_O_7_) and oxalic acid (C_2_H_2_O_4_) (Chaboud, 1983; Collins & Reilly, 1968), are recognised as major compounds in the literature, but do not accurately reflect the composition of the root-derived exudates analysed in this study. For measuring mineralisation kinetics of the latter two organic acids, soils were buffered by a citrate buffer (0.1 M C_6_H_8_O_7_, 0.1 M C_6_H_5_Na_3_O_7_ · 2H_2_O) to keep the pH 5.6 of the soil constant. All reagents used in this study were analytical grade and purchased from Sigma Aldrich (Merck, Darmstadt, Germany), unless otherwise stated.

Exudates and mucilage of maize (*Zea mays* B73 wild type) were collected in the same field experiment as described above (Vetterlein et al., 2021). Exudates were collected in the early vegetation period during leaf development (BBCH 14), and in the late vegetation period at the first ripening stage (BBCH 83) (Lancashire et al., 1991) by a soil hydroponic-hybrid approach as described in Santangeli et al. (2024). Briefly, soil-grown plants were grown in perforated soil columns in the field which allowed excavation of the entire intact maize plants. The root system was gently rinsed with tap water for 30 min to remove soil particles. Root exudates were then collected hydroponically for one hour in 0.5 L and 7.5 L of deionised water at BBCH 14 and BBCH 83, respectively. Finally, the solution was filtered (0.2 µm, Cellulose acetate OE 66, Whatman, UK) to remove all root debris, to capture only the soluble exudate fraction. The filtered exudates were divided into aliquots, frozen at -20°C and stored at -80°C until analysis. Exudate C contents were determined on a Shimadzu TOC-5050 from freeze-dried subsamples.

Immediately before mineralisation measurements, the remaining freeze-dried material was dissolved in ultra-pure H_2_0 to obtain the desired C concentration.

Mucilage collection followed the procedure in Ahmed et al. (2015), with slight modifications. Briefly, nodal roots were carefully detached from the maize root systems at the end of tassel emergence (BBCH 59) and cleaned from soil with deionized water. Nodal roots with intact tips were submerged in deionized water for 24h to allow mucilage hydration. Subsequently, the excess water was discarded through a fine sieve and hydrated mucilage was aspirated using syringes and frozen at -20°C until use. The C and N content of the mucilage was analysed by combustion using a C/N elemental analyser (Thermo Flash EA 2000, Fisher Scientific). The remaining material was dissolved in ultra-pure H_2_0 to obtain the desired C concentrations.

### Measurements of microbial substrate utilization

Microbial utilisation of single rhizodeposit components, maize exudates and mucilage (hereinafter referred to as ’substrates’) was measured by an electrolytic O_2_ microcompensation apparatus as described in Scheu (1992). To mimic the decrease of rhizodeposits and the resulting lower C supply to the soil microorganisms with increasing distance from the root surface, the individual substrates were diluted with ultrapure H_2_O according to their C content to form a concentration gradient of 1600, 400, 200, 100 and 40 μg C g^-1^ soil dry wt. For mucilage, mineralisation of four additional concentrations of 1200, 1000, 800 and 600 μg C g^-1^ soil were measured. For early and late season exudates the concentration step of 200 μg C g^-1^ soil was omitted.

Samples of field soil, equivalent to 3 g dry weight, were adjusted to 60% of their maximum water holding capacity (corresponding to a gravimetric water content of 30%) to ensure uniform water content. The samples were either amended with a substrate or without, i.e. microbial basal respiration. Thus, equivalent amounts of water have been used to add different quantities of C. Oxygen consumption rates at 22°C were monitored every 15 minutes for the next 24-48 hours with three replicates, each. In order to assess the C mineralisation of the substrates, the microbial O_2_ consumption was converted into respired CO_2_-C according to the ideal gas law and assuming a respiratory quotient of 1. In addition, the extent to which the microbial mineralisation of the added C substrates was limited by the availability of N and P was determined by amending the soils in a factorial combination with solutions of (NH_4_)_2_SO_4_ or/and KH_2_PO_4_, corresponding to an optimal microbial C:N:P stoichiometry of 10:2:1 (Griffiths et al., 2012).

### Calculations of substrate mineralisation kinetics

In order to analyse the mineralisation of rhizodeposits, microbial kinetic substrate-induced respiration (KSIR) was measured from the respiration curves (Blagodatsky et al., 2000) (Figure 1). First, the microbial basal respiration rate [μg CO_2_-C g^−1^ h^−1^] was determined in soil without substrate addition. The addition of a substrate causes an immediate significant upregulation of microbial respiration above the level of basal respiration prior to the onset of microbial growth. During this lag-period, the MIRR (Anderson & Domsch, 1978), was calculated as the mean of the three lowest respiration measurements. Microbial biomass (Cmic) was calculated from the mean MIRR value after addition of 1600 µg C g^-1^ glucose according to the equation (1) (Beck et al., 1997):

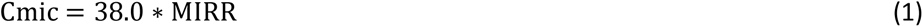

**Figure 1.**
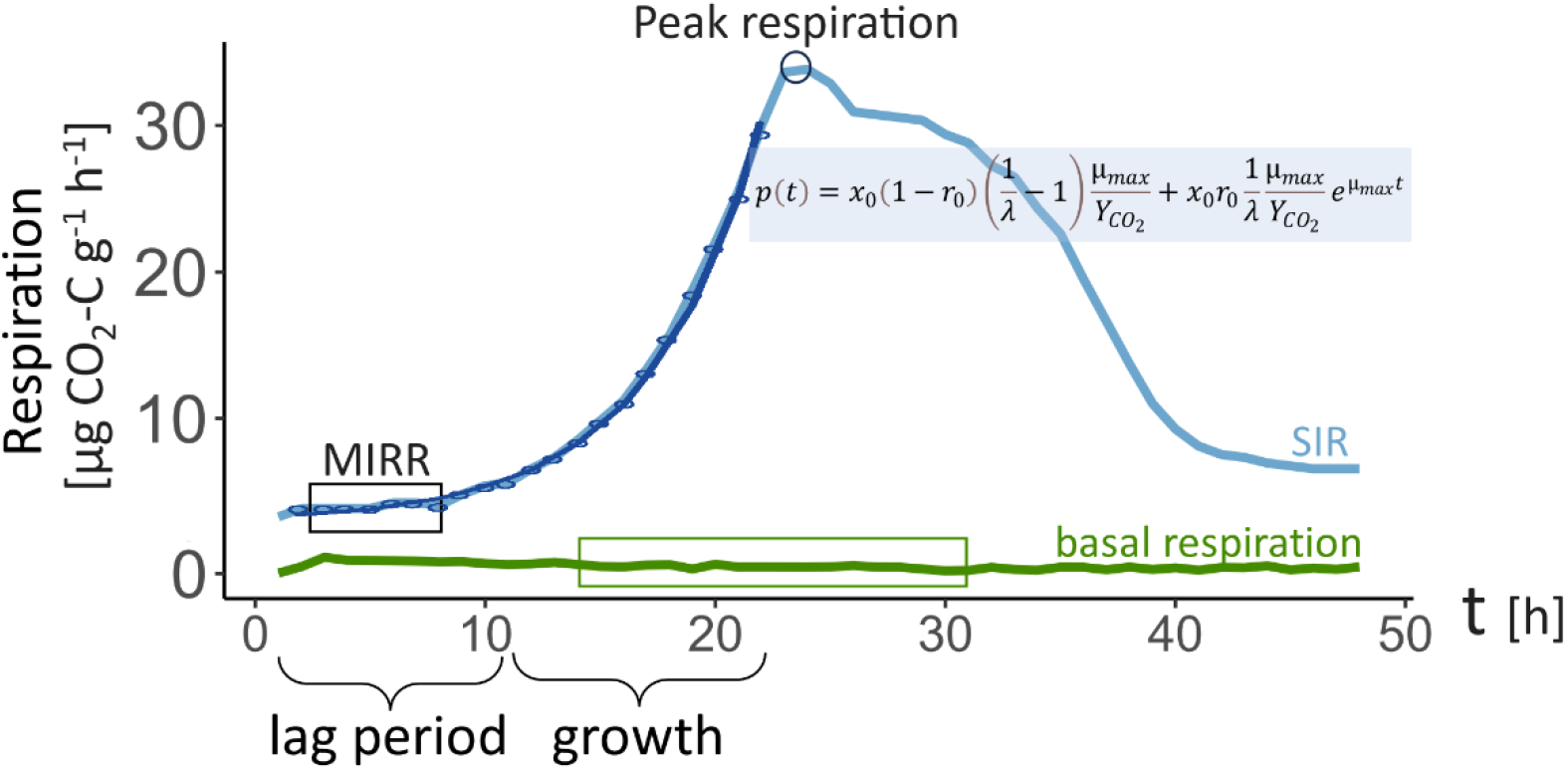
Explanatory graph for definition of respiration measurements by microbial growth behaviour. MIRR is the maximal initial respiratory response, calculated as the mean of the three lowest lag period measurements. SIR is substrate induced respiration. With the kinetic model by Wutzler et al. (2012) the respiration curve is fitted to the respiration rates and used to calculate microbial parameters.

The specific respiration (qCO_2)_ could then be calculated as the ratio between basal respiration and Cmic [μg CO_2_-C μg Cmic-C^-1^].

Substrate supply in excess to microbial demand allowed for unrestricted exponential growth of microorganisms, but growth of microbial biomass decreased with the stepwise reduction of the C supply until no further growth occurred. The time span of the lag-period until the exponential increase of respiration during microbial growth and its decrease during the microbial depletion of the substrate were covered by the measurements. The differences in substrate application rates enabled the determination of the C-threshold, which reflects the transition from maintenance to active growth of the microorganisms (Panikov, 1995). Therefore, the SIR approach allows the calculation of the following microbial mineralisation characteristics: i) the C-threshold needed to initiate microbial growth, which was determined according to Anderson and Domsch (1985) as the concentration that merely triggered an increase in respiration, which remained constant for several hours before decreasing, but at which the C supply was insufficient for microbial growth, ii) the peak respiration was set as the first peak in microbial growth, typically occurring between 10-30 h after the addition of the substrate. Additionally, iii) the maximum specific growth rate (μ_max_) and iv) the active microbial fraction were estimated using KSIR analysis. The KSIR analysis was based on the respiration measurements in response to the C concentration that gave the highest peak respiration for each substrate (Blagodatsky et al., 2000). Following the model presented in Wutzler et al. (2012), the measured respiration rates were described by the equation (2) and best-fit parameters were estimated.

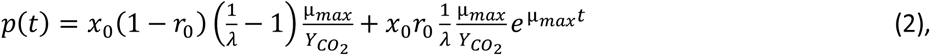

where p(t) is the respiration rate at time t, expressed as respired C per time, µ_max_ is the maximum specific growth rate, r_0_ is the initial physiological state, x_0_ is the microbial biomass at the moment of substrate addition. As suggested in Wutzler et al. (2012), (2) was used as a three-parameter equation, accepting the following assumptions: first, λ was assumed to be a basic stoichiometric constant of 0.9 during unrestricted growth (Akimenko et al., 1983). Secondly, Y_CO2_ = Y / (1 - Y) can be assumed to be a constant of 1.5 during unlimited growth (Blagodatsky et al., 2000). These values were subsequently used to calculate the lag-period (t_lag_) as described in Baranyi and Pin (1999). T_lag_ is inversely proportional to µ_max_ and depends on r_0_. It was determined using equation (3):

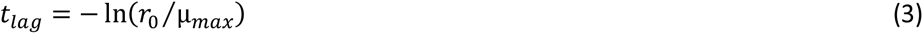

For each substrate, a Michaelis-Menten model curve was fitted to the MIRR values of the different concentration steps, relating the added substrate concentration to the initial microbial respiration rate. This model was used to calculate the maximum respiration rate (V_max_), which indicates the theoretical microbial peak CO_2_ production during MIRR for a specific substrate, and the Michaelis-Menten constant (K_m_), describes the affinity of microorganisms to mineralize the given substrate, with low values indicating high microbial substrate mineralisation at low substrate concentrations. The turnover time (T_t_) of the applied substrate is the total time required by the microbial community to metabolise the applied substrate at K_m_ plus the concentration of soil indigenous C, measured as EOC. This estimation was made using equation (4)(Blagodatskaya et al., 2009):

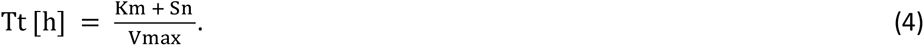

Two-way ANOVA was used to analyse differences between MIRR and peak respiration dependent on substrate and nutrient limitation, with Tukey’s test for post-hoc pairwise comparisons. Calculations and visualisation of results were performed using R statistical software (version 4.2.1, 2022-06-23; “Funny-Looking Kid”). The following R packages were utilized for various aspects of the analysis: ’ggplot2’ for data visualization (Wickham et al., 2018), ’dplyr’ to arrange the data (Wickham, 2020), ’reshape2’ for melting data (Wickham, 2007), ’nlstools’ for Michaelis-Menten analysis (Baty et al., 2015), and ’twKinresp’ for KSIR (Macholdt, 2017).

## Results

### Microbial mineralisation of rhizodeposits at varying substrate concentrations

The loamy agricultural soil had a microbial biomass of 158 μg Cmic-C g⁻¹ with a basal respiration of 0.18 μg CO_2_-C g^-1^ h^-1^. Accordingly, the specific respiration (qCO_2_) was 1.14 μg CO_2_-C g^-1^ Cmic-C. In comparison, the maize mucilage contained 357 µg C g^-1^ and 16.2 µg N g^-1^ (C:N ratio = 22), while early season exudates (BBCH 14) and late season exudates (BBCH 83) had C contents of 10.9 and 12.8 µg C ml^-1^, respectively.

Microbial respiration was increased well above basal respiration after the addition of sugars and amino acids, with a characteristic lag-period lasting until the onset of exponential microbial respiration when the substrate supply exceeded the substrate-specific C-threshold for microbial growth (Figure 1, **Fehler! Verweisquelle konnte nicht gefunden werden.**). The extent of this increased microbial respiration was dependent on the concentration of substrate added. In response to the sugars applied, respiration increased 12-fold above basal maintenance levels during the first 10 h to a MIRR of 2.25 μg CO_2_-C g^-1^ h^-1^ after the addition of 1600 µg C g^-1^ glucose or sucrose, and 9-fold to 1.65 μg CO_2_-C g^-1^ h^-1^ in response to 1600 µg C g^-1^ arabinose (Table 1, Figure 2). The lag-phase with increased respiration turned into exponential respiration at the onset of microbial growth. The C-threshold for microbial growth on glucose and sucrose was 100 µg C g^-1^. For arabinose, the threshold C concentration required to initiate microbial growth was lower with 40 µg C g^-1^ (**Fehler! Verweisquelle konnte nicht gefunden werden.**). Microbial respiration peaked 22 hours after the addition of glucose. Higher substrate levels resulted in higher rates of peak respiration, indicating higher microbial growth. Below the C-threshold for microbial growth however, the initial upregulation of respiration after glucose addition dropped off without exhibiting the typical exponential phase seen at higher substrate levels (Figure 2). At peak respiration, approximately 0.01% of the total amount of added sugar-C was respired per hour (Table 1, Figure 2). The total (cumulative) CO_2_-C respired from sugars at peak respiration amounted to between 7-9% of the added C. Peak respiration following arabinose addition at 27 h was delayed by 5 and 7 h longer compared to glucose and sucrose, respectively (Table 1, Figure 2).

**Figure 2.**
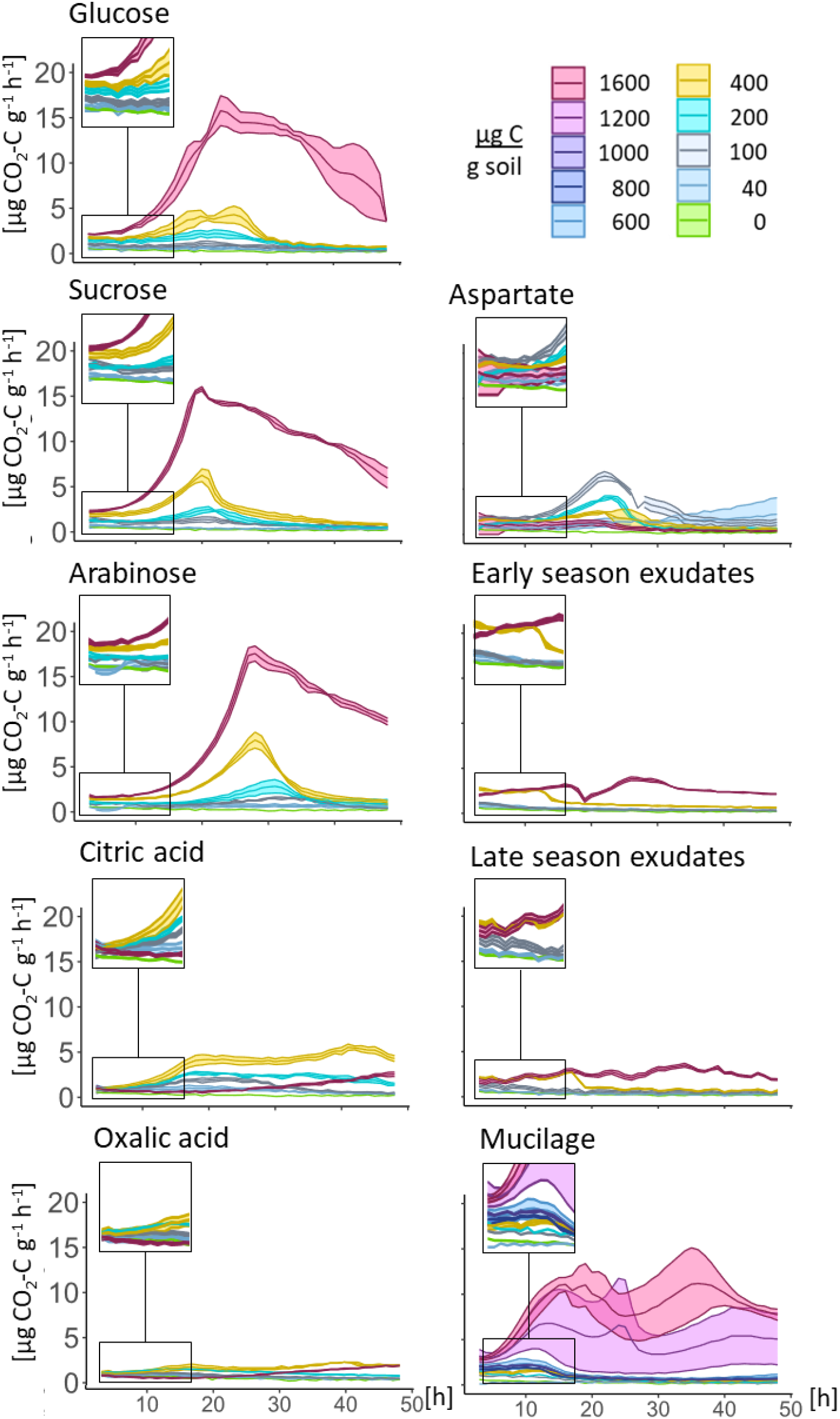
Microbial growth in response to different root derived substrates: glucose, sucrose, arabinose, aspartate, critic and oxalic acid, mucilage, early and late exudates. Each curve represents the mean of three replicates +/- standard error. The lag period is enlarged in the top left corner of each graph, with the scale of the enlargement indicated by rectangles.

**Table 1.** Microbial mineralisation response parameters to the different substrates without and with nutrient supplementation. Only the values for each substrate that correspond to the C concentration resulting in the highest peak respiration are displayed. Results are the average of 3 replicates.

| Substrate | + | MIRR |  |  | Peakrespiration |  | cumulative CO<sub>2</sub><br>respiration until peak |  | Time of Peak<br>respiration |
| --- | --- | --- | --- | --- | --- | --- | --- | --- | --- |
|  |  | µg C g<sup>-1</sup> | µg CO<sub>2</sub>-C g<sup>-1</sup>h<sup>-1</sup> | SE | µg CO<sub>2</sub>-C g<sup>-1</sup>h<sup>-1</sup> | SE | µg CO<sub>2</sub>-C g<sup>-1</sup> | SE | [h] |
| Glucose |  | 1600 | 2.23 | 0.09 | 16.20 | 1.31 | 127.23 | 14.73 | 22.00 |
|  | P | 1600 | 2.09 | 0.06 | 19.87 | 0.39 | 152.64 | 7.37 | 23.33 |
|  | N | 1600 | 2.41 | 0.43 | 22.84 | 1.15 | 188.98 | 8.82 | 22.33 |
|  | NP | 1600 | 1.80 | 0.24 | 21.15 | 1.46 | 199.02 | 12.69 | 22.67 |
| Sucrose |  | 1600 | 2.28 | 0.11 | 15.92 | 0.15 | 112.04 | 2.50 | 19.67 |
|  | P | 1600 | 2.32 | 0.18 | 17.31 | 0.81 | 110.83 | 4.05 | 19.33 |
|  | N | 1600 | 2.25 | 0.30 | 28.35 | 1.02 | 218.82 | 16.11 | 20.67 |
|  | NP | 1600 | 2.17 | 0.68 | 19.67 | 5.57 | 195.26 | 7.32 | 20.50 |
| Arabinose |  | 1600 | 1.65 | 0.07 | 17.55 | 0.83 | 146.44 | 6.32 | 27.00 |
|  | P | 1600 | 1.41 | 0.20 | 11.56 | 0.21 | 82.97 | 6.66 | 24.00 |
|  | N | 1600 | 1.87 | 0.07 | 16.89 | 0.31 | 122.63 | 1.99 | 24.00 |
|  | NP | 1600 | 1.22 | 0.91 | 10.73 | 2.87 | 99.34 | 27.34 | 26.00 |
| CiAcid |  | 400 | 0.98 | 0.13 | 4.33 | 0.44 | 39.49 | 3.69 | 19.67 |
| OxAcid |  | 400 | 1.04 | 0.12 | 1.87 | 0.16 | 19.42 | 2.66 | 15.33 |
| Aspartate |  | 100 | 1.77 | 0.13 | 6.47 | 0.46 | 60.85 | 5.02 | 21.00 |
|  | P | 100 | 1.06 | 0.45 | 4.48 | 1.77 | 56.44 | 4.52 | 23.00 |
| early |  |  |  |  |  |  |  |  |  |
| Exudates |  | 1600 | 2.05 | 0.04 | 3.88 | 0.19 | 68.19 | 3.23 | 26.33 |
| late |  |  |  |  |  |  |  |  |  |
| Exudates |  | 1600 | 1.65 | 0.24 | 3.50 | 0.21 | 81.34 | 6.85 | 34.00 |
| Mucilage |  | 1600 | 2.92 | 0.18 | 15.13 | 1.58 | 143.55 | 20.75 | 19.25 |

SE: Standard error of the mean, MIRR: maximum initial respiratory response

The C-thresholds for microbial growth on citric and oxalic acid were 40 and 100 µg C g^-1^, respectively. The addition of organic acids to buffered soils at concentrations above 400 µg C g^-1^ inhibited microbial activity, with growth starting only after 20-21 h. The highest peak respiration of 4.3 µg CO_2_-C g^-1^ h^-1^ was observed after the addition of 400 µg C g^-1^ citric acid, which is the same magnitude as the respiration peak in response to an addition of sugars at 400 µg C g^-1^. The cumulative respired CO_2_ at peak respiration after the addition of citric acid amounted to 10% of the total added C, while for oxalic acid it was 5% of total added C.

The soil microorganisms mineralised the amino acid aspartate more efficiently than sugars. The C-threshold for microbial growth was 40 µg C g^-1^ aspartate. The addition of 100 µg C g^-1^ aspartate resulted in a peak respiration rate of 6.5 µg CO_2_-C g^-1^ h^-1^. This means that 0.07% of the substrate_C was respired per hour at peak respiration, leading to a cumulative C respiration of 61% of the added C until peak respiration, 21 h after substrate addition (Table 1). However, aspartate concentrations above 400 µg C g^-1^ inhibited microbial growth.

Patterns of microbial mineralisation of root-derived C sources such as exudates and mucilage were more complex. The threshold for microbial growth was only exceeded in response to 400 and 100 µg C g^-1^ of early and late season exudates, respectively (**Fehler! Verweisquelle konnte nicht gefunden werden.**). However, even when the C-threshold for exudates was exceeded, the microbial respiration did not enter an exponential growth phase as with the other substrates, but merely a steady increase was observed (Figure 2). This led to a peak respiration of about 3-4 µg CO_2_-C g^-1^ h^-1^ in response to exudates at 1600 µg C g^-1^ (Table 1). For mucilage, the C-threshold for microbial growth was at 1000 µg C g^-1^ (**Fehler! Verweisquelle konnte nicht gefunden werden.**), leading to a peak respiration almost as high as for sugars (up to 13.4 µg CO_2_-C g^-1^ h^-1^) (Table 1). Early season exudates and mucilage caused two respiratory growth peaks at this highest C concentration (Figure 2). The first peak was observed 24 h after substrate addition as for the other substrates, the second peak 30-35 h after substrate addition. Microbial respiration at the highest concentrations of late-season exudates decreased very slowly after the initial peak, without showing the typical decline in microbial activity after one day. The cumulative respired CO_2_-C at peak respiration was about 4% of the added C in early and late season exudates, and 9% of the added C for mucilage (Table 1).

Overall, the microbial mineralisation of rhizodeposits showed more complex patterns as compared to sugars, with multiple growth peaks and generally requiring higher C concentrations to initiate microbial growth.

### Microbial growth kinetics

The kinetic variables obtained from the model fitting of the respiration curves showed high similarity among the sugars (**Fehler! Verweisquelle konnte nicht gefunden werden.**). The specific growth rate (µ_max_) of the microbial community on the monosaccharide sugars glucose (µ_max_ = 0.21) and arabinose (µ_max_ = 0.20) was slightly lower than the specific growth rate on the disaccharide sucrose with 0.26 (Figure 4). Microbial growth on aspartate also showed a high µ_max_ (0.27), but with a longer lag-period compared to sugars, and aspartate stimulated a very low active fraction of Cmic (Figure 5). In response to mucilage, microorganisms showed a relatively low µ_max_ of 0.18. Mucilage stood out due to its notably brief lag time of 58 min and a more than tenfold higher active fraction of Cmic than any of the other tested substrates (Figure 5). Due to the absence of a distinct growth phase, the respiration curves of organic acids and exudates could not be fitted to assess the lag period, µ_max_, and the active fraction of microbial biomass according to the Wutzler model.

### Michaelis-Menten kinetics

The relationship between the quantity of substrate added and MIRR, shown by Michaelis-Menten kinetics (Figure 3), confirms significant variation in the mineralisation of different rhizodeposit substrates. Across a range of substrates, respiration showed a sharp increase up to roughly 250 µg C g^-1^ before reaching a plateau near V_max_. As expected, microorganisms showed a high affinity to sugars and exudates with low K_m_ values at around 100 µg C g^-1^ (Figure 4, **Fehler! Verweisquelle konnte nicht gefunden werden.**). Glucose, sucrose and early-season exudates had a high V_max_ value of around 2.5 µg CO_2_-C g^-1^ h^-1^ and T_t_ about 3h, whereas the late-season exudates and arabinose had lower V_max_ values, falling below 2 µg CO_2_-C g^-1^ h^-1^ and T_t_ about 4h. The Michaelis-Menten model was successful in fitting the mineralisation of the easily decomposable sugars and exudates, but no K_m_ values could be determined for aspartate, the organic acids and mucilage (Figure 3).

**Figure 3.**
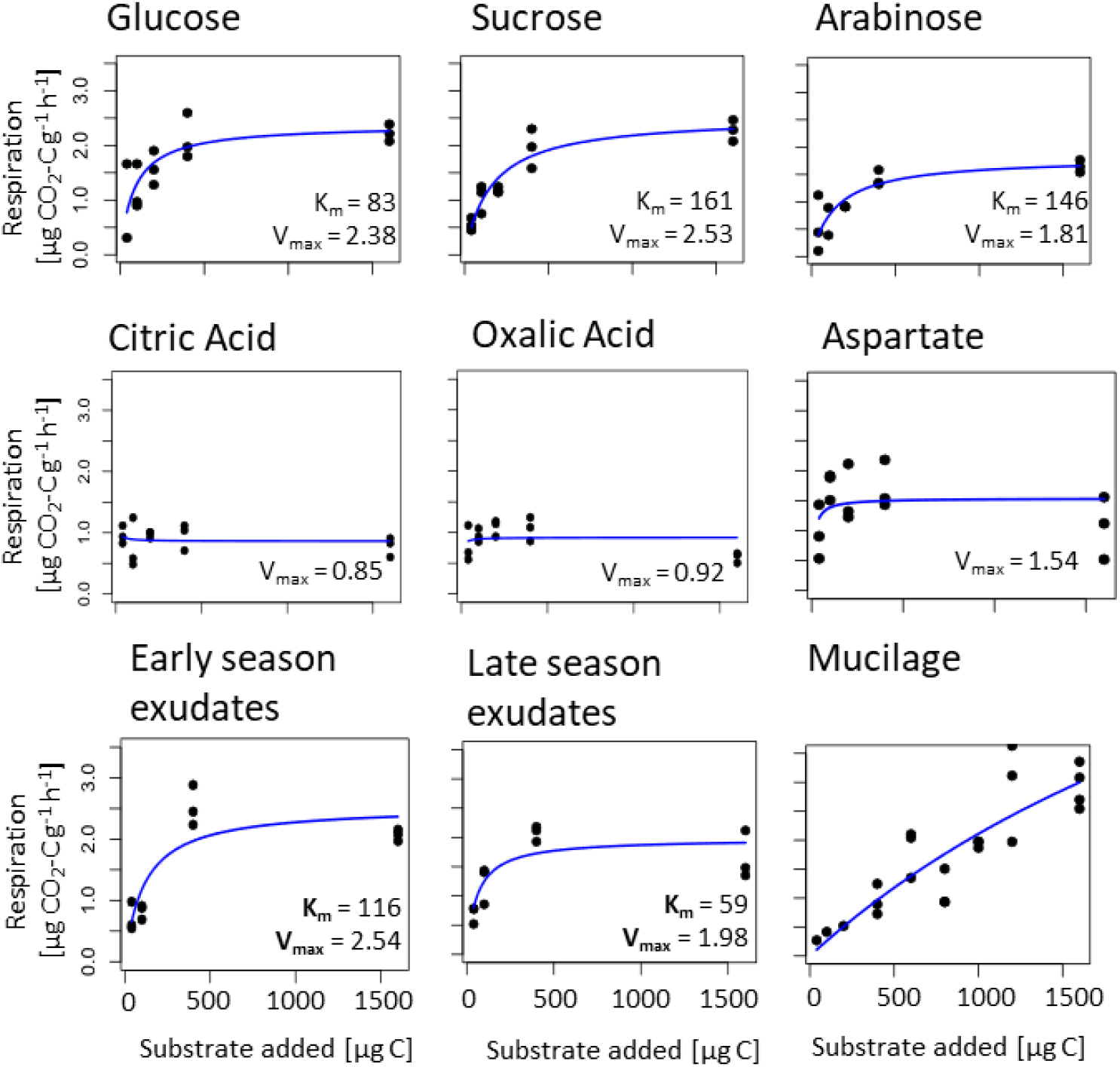
Michaelis-Menten kinetics, Model of MIRR [µg CO_2_-C g^-1^ h^-1^] to a gradient of substrate concentrations [µg C] (glucose, sucrose, arabinose, aspartate, citric acid, oxalic acid, early season exudates, late season exudates and mucilage), K_m_ and V_max_ values are displayed if significant.

The model was more robust for calculations of V_max_ (**Fehler! Verweisquelle konnte nicht gefunden werden.**), except for mucilage, where the initial upregulation of microbial respiration (MIRR) increased linear with the added amount (MIRR = 0.30 + 1.74 * µg C mucilage g⁻¹, R²= 0.75) without reaching a plateau at the highest concentration of 1500 µg C g^-1^ soil (Figure 2).

### Effect of nutrient addition on substrate-induced respiration response

Microbial growth in soil is mainly limited by C and was therefore strongly stimulated by the C-rich rhizodeposits. The addition of N and P on top of the C input by glucose or sucrose showed a significant increase in peak respiration in response to the combination of C and N, demonstrating a strong shift in nutrient limitation of microbial growth to N (Figure 5). However, a microbial co-limitation of N and P was indicated by the amplification of peak respiration and a decrease of the C-threshold required for microbial growth when both nutrients together were added with glucose (**Fehler! Verweisquelle konnte nicht gefunden werden.**, Table 1, **Fehler! Verweisquelle konnte nicht gefunden werden.**). By the addition of N, P or both to glucose, microbial growth was initiated already at 40% of the threshold concentration required without it (**Fehler! Verweisquelle konnte nicht gefunden werden.**). Notably, in response to sucrose a reduction of the C-threshold was only observed with N-supplementation. In contrast, nutrient addition to arabinose raised the C-threshold for the transition from maintenance to microbial growth, corresponding to an increment to the next substrate concentration level, e.g. from 40 to 100 µg C g^-1^ with the addition of nutrients (**Fehler! Verweisquelle konnte nicht gefunden werden.**).

**Figure 4.**
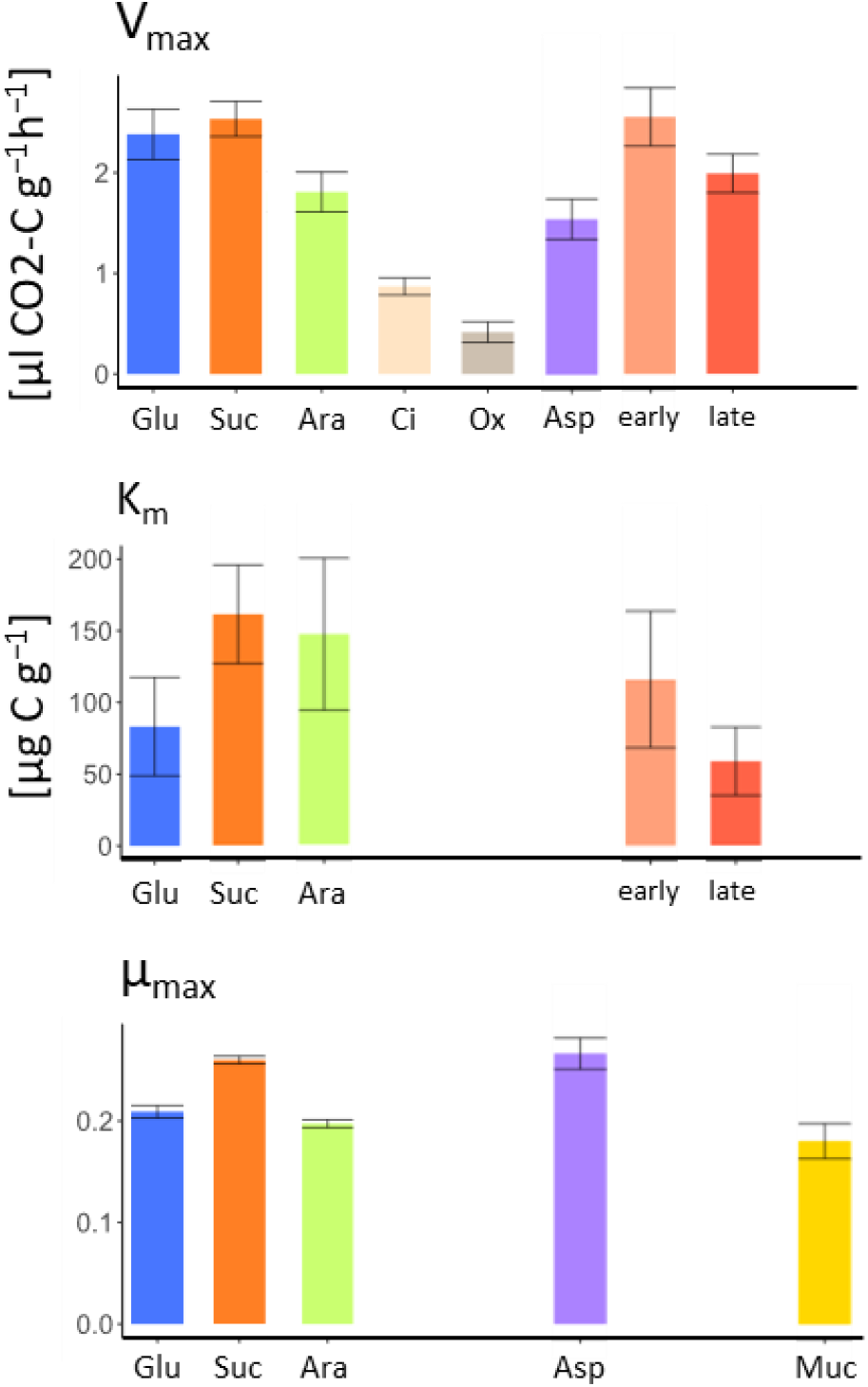
Community properties in response to different substrates. The bar graphs show the Michaelis-Menten model-based parameters for V_max_ and K_m_, and the KSIR model-based parameters for µ_max_, with standard error of V_max_ and K_m_ and standard deviation of µ_max_. Substrates include Glu (Glucose), Suc (Sucrose), Ara (Arabinose), Ci (Citric acid), Ox (Oxalic acid), Asp (Aspartate), early and late (early and late season exudates) and Muc (Mucilage).

**Figure 5.**
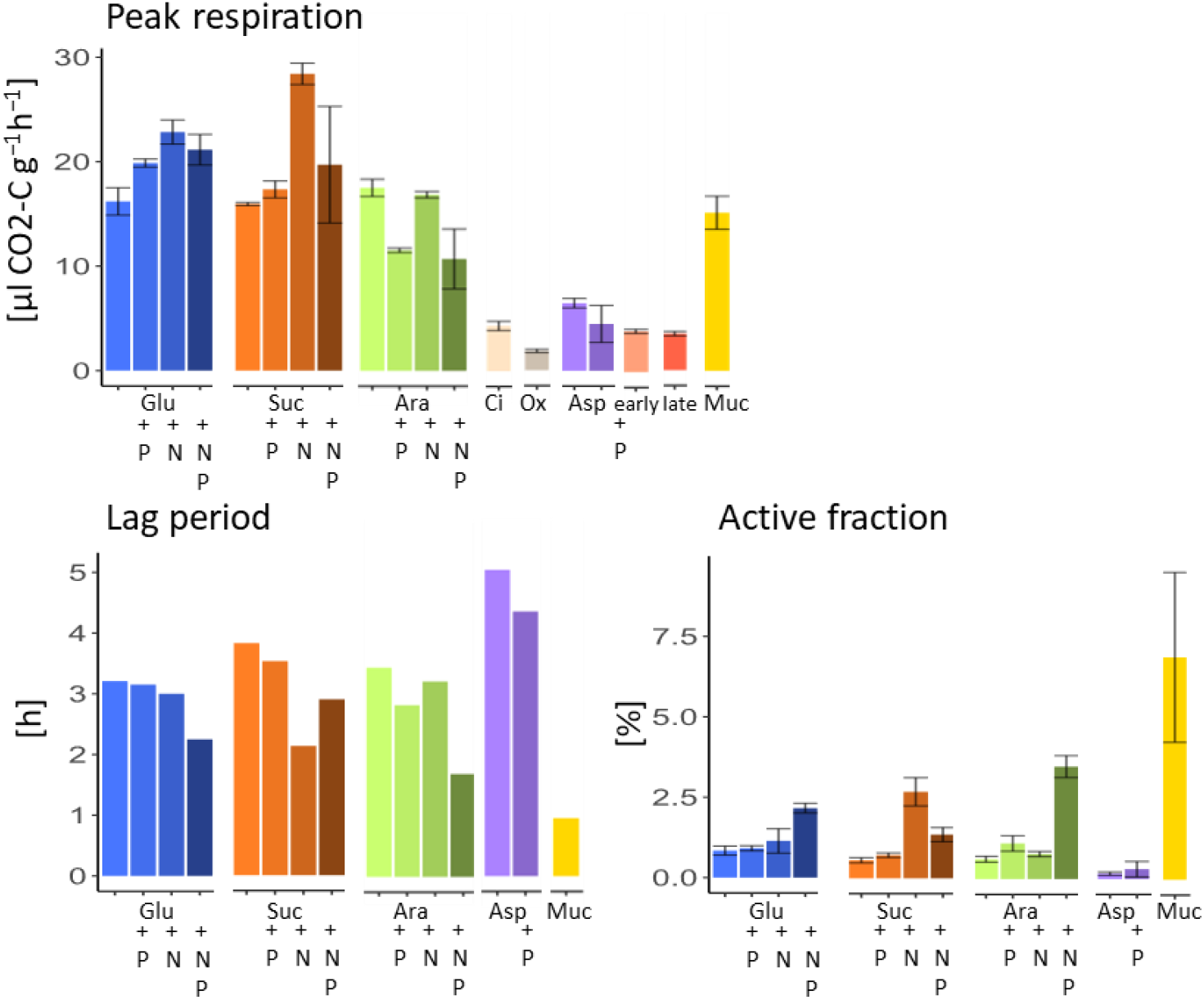
Parameter for microbial growth on different substrates with and without nutrient limitations. Substrates include 1600 µg C g^-1^ Glu (Glucose), 1600 µg C g^-1^ Suc (Sucrose), 1600 µg C g^-1^ Ara (Arabinose), 400 µg C g^-1^ Ci (Citric acid), 400 µg C g^-1^ Ox (Oxalic acid), 100 µg C g^-1^ Asp (Aspartate), 1600 µg C g^-1^ early and late (early and late season exudates) and 1600 µg C g^-1^ Muc (Mucilage). The bar graph shows the measured value of peak respiration, and the KSIR model-based parameters for lag-period [h] before exponential growth, and active fraction (the active part of the total biomass), n=3, with standard error of peak respiration and standard deviation of difference for active fraction.

Overall, the lag-period was reduced by 50-75% and the active fraction of the microbial biomass increased 2.5- to 6-fold with the addition of both N and P (Figure 5, **Fehler! Verweisquelle konnte nicht gefunden werden.**).

## Discussion

Rhizodeposits are a diverse mixture of organic compounds released by plant roots that stimulate microbial activity and influence the physical and chemical properties of the rhizosphere (Carminati, 2013; Nazari, 2021). However, studies investigating microbial responses to substrate inputs into the soil often use glucose as a model substrate (Anderson & Domsch, 1985; Reischke et al., 2015; Sawada et al., 2008; Stenström et al., 2001) and sometimes other single components of rhizodeposits such as organic acids (Van Hees, 2002). We observed a typical dynamic of the microbial degradation of sugars with a clear lag period before the exponential growth takes place at peak respiration at 22h. The observed patterns confirm a strong sugar affinity of the microbes and high mineralisation rates. The aldopentose arabinose however, was utilised with slower turnover by fewer active microbes. Apparently, the bulk soil microbiome was not well adapted for a rapid metabolisation of arabinose. Although arabinose is a common constituent of rhizodeposits, it is far less abundant than glucose (Chaboud, 1983; Nazari et al., 2020). The difference to the other sugars may be due to potentially less abundant or fewer activated transporters involved in arabinose mineralisation compared to glucose transporters (Mayer, 2005; Ryu, 2018).

From a stoichiometric perspective, amino acids provide both C and N for optimal microbial growth. Accordingly, microbial growth was significantly increased at 100 µg C g^-1^ of aspartate, but elevated concentrations of aspartate did inhibit microbial growth. Amino acid uptake may be also hampered by physiological limitations of non-adapted microbial bulk soil communities (Koskella, 2015), as the numbers and activity of specific ABC transporters restricts the microbial uptake of amino acids (Hosie & Poole, 2001).

The adaptation of different microbial taxa to degrade individual substrates and the complexity of substrates may explain the temporal variation in the microbial mineralisation of exudates and mucilage in our study. Rhizodeposits could trigger microbial growth through the provision of readily available C substrates, but they also contain chemical inhibitors of microbial growth (Nazari, 2021; Oburger et al., 2011; Wiesenbauer et al., 2023). Two distinct growth peaks in response to early season exudates and to mucilage indicate interactions between taxa that utilize different C sources at different times, and potentially the influence of compounds that delay substrate utilisation (Badri, 2009; Baetz, 2014). In particular the late season exudates did exhibit multiple small respiration peaks, some of which may be superimposed and therefore not visible, but still contributing to the rugged pattern of microbial growth on late season exudates (Figure 2).

The assembly of the rhizosphere microbiome starts at the root tip and was shown to change along the root axis towards older root sections (Dupuy & Silk, 2016; Rüger et al., 2021). The bulk soil microbial community is mostly dormant and its small active part dominated by K-strategists, strictly adapted to resource-limited conditions (Soler-Bistue et al., 2023). As the root extends into the bulk soil, the secretion of mucilage at root tips activates the dormant bulk soil microbiota and creates a first temporally and spatially restricted hotspot of microbial activity (Kuzyakov & Blagodatskaya, 2015). The brief lag phase following mucilage addition with a low µ_max_ indicates the growth of a significant proportion of K-strategists (Couso et al., 2023). Due to their lower energy requirements these K-strategists are capable of immediate growth upon encountering a mucilage-coated root tip. However, these K-strategists do not reproduce as efficiently as r-strategists, reflected by their relatively low μ_max_ for the immediate microbial growth response to mucilage supplementation. In addition, soluble exudates are released behind the root tip into the soil matrix, where mucilage has already induced the growth of a significant proportion of the microbial biomass. The majority of maize exudates is composed of sugars (Santangeli et al. (2024), and therefore one would intuitively expect their rapid microbial metabolisation, and a replacement of K-strategists by r-strategists with faster turnover rates. Interestingly, after the typical initial up-regulation of microbial metabolism in response to root exudates, the expected exponential microbial growth was not realised. Secondary metabolites in exudates repeatedly have been found to restrict the growth of rhizosphere microbiota (el Zahar Haichar, 2014; Khashi u Rahman, 2019; Sasse et al., 2018). One likely reason is that plants hinder the rapid degradation of substances with functional potential by involving antibiotics and microbial inhibitors in rhizodeposition (Bais et al., 2006; Hawes, 2000). The rather slow microbial growth rate on root exudates fit well with a recent study by (Zhalnina, 2018), who inferred from a codon-usage bias in members of the rhizosphere microbiome of the grass *Avena*, and later confirmed by culturing studies, that the majority of rhizosphere bacteria had much longer generation times than would be expected from fast-growing r-strategists.

There is further a spatial component in addition to the temporal utilization of rhizodeposits. The C supply to microorganisms by rhizodeposits is decreasing rapidly with distance from the root surface (Alphei et al., 1996), this was mimicked by the stepwise addition of lower concentrations of rhizodeposits to soil. When confronted with a limiting C-source, microorganisms make strategic choices, as the uptake of substrate and its conversion into storage compounds requires less energy than the synthesis of cellular structural compounds during growth (Sawada et al., 2008). This results in a critical threshold that differentiates between the microbial use of C for storage, or its use for growth when a critical C-threshold is exceeded (Bremer & Kuikman, 1994). As the microbial biomass increases, the required C-threshold also tends to increase as well (Reischke et al., 2015) and lies approximately at 30% of Cmic, according to Anderson and Domsch (1985). Therefore, any C-threshold obtained should be normalised by the soil’s Cmic content to make studies on different soils comparable. Our data show a Cmic-specific threshold between 25-63% to trigger microbial growth in response to sugars, aligning with previous studies (Anderson & Domsch, 1985; Reischke et al., 2015; Sawada et al., 2008). In addition, the C-threshold for microbial growth varies with the type of C-substrate. The C-thresholds for late season exudates were not different from those for sugars, amino acids, and organic acids. This corresponds to the findings of Santangeli et al. (2024) reporting that late-season exudates consist to over 50% of soluble carbohydrates. On the other hand, the C-thresholds for microbial growth in response to early season exudates and mucilage were an order of magnitude higher, at 2.53 and 6.33 µg C µg Cmic-C^-1^ respectively, indicating that these rhizodeposits were less supportive for microbial growth. It has been shown that the composition of the rhizosphere microbiome is closely linked to the bouquet of exudates that plants release in order to recruit members of the bulk soil microbial community (Zhalnina, 2018). However, only plants in their growth phase (e.g. BBCH 14) will benefit most from any plant-protective or growth-promoting microbial traits, while after flowering all resources are redirected to seed formation (e.g. BBCH 83) (Keith et al., 1986; Swinnen et al., 1994). The decrease of secondary phenolic compounds in late season exudates at ripening (BBCH83) of maize (Santangeli et al., 2024) may be an indication of the switch from microbiome selection to general microbial growth promotion by rhizodeposits. When relating substrate concentration to the initial microbial response using Michaelis-Menten kinetics, microbial growth on early season exudates showed a more effective substrate turnover (lower T_t_) due to the activation of a larger fraction of the microbial community (higher V_max_), but with reduced affinity to the substrate, leading to an increased C-threshold for growth on early season exudates. Microbial growth dynamics on early season exudates thus resembled the pattern of the polysaccharide sucrose (Fig. 3). Late season exudates in contrast supported the growth of microorganisms with a lower K_m_ and a longer T_t_ for late season exudates.

Also for stoichiometric reasons, it may not be in the interest of plants that their rhizodeposits fuel microbial growth too rapidly. The C-supply from roots lifts the C-limitation of the soil microorganisms, whose growth then becomes nutrient-limited. Unfortunately, the amount of rhizodeposits available was not sufficient to measure the nutrient limitation of microbial growth. However, the increased peak respiration after amending sugars with N and P demonstrates the hypothesized switch towards strong N-limitation of microbial growth (Figure 5), leading to strong nutrient competition with plants (Hodge et al., 2000). The reduced lag-phase and the strongly increased fraction of microorganisms activated by the substrate, especially when supplied with both N and P (Figure 5), give further proof of how strongly the microbial mineralisation of these substrates was nutrient-limited. For mucilage, where we could obtain measurements of C and N contents, its C:N ratio of 22 was much wider than the C:N of 5 applied to soils. Mucilage, despite much lower N-availability, exhibited the shortest lag period, and activated the largest fraction of the soil microbial biomass (Figure 5). Thus not only nutrients, but additional components within the mucilage likely accelerated the rapid activation of a high fraction of the microbial biomass, perhaps aided by enzymes that facilitate the breakdown of polysaccharides for microbial use and nutrient uptake (Pozzo et al., 2018), or by the recruitment of special microorganisms, like those containing glycosyl hydrolases to degrade polysaccharides (Amicucci et al., 2019).

## Conclusion

The microbial mineralisation of rhizodeposits showed more complex temporal patterns than investigations on individual model substrates would suggest. A high C-concentration of 1000 µg C mucilage g^-1^ was needed to stimulate exponential microbial growth, but then mucilage stimulated two growth peaks. Contrary and unexpected was the low microbial growth on exudates, and the remarkable differences in the microbial use of early and late season exudates. Overall, the initiation of microbial growth required surpassing a critical C-threshold, while lower C availability primarily stimulated microbial respiration. Accordingly, at greater distance to the root surface, where the C-supply via rhizodeposition falls below this critical threshold, the rhizosphere microorganisms preferentially respire the assimilated C without converting it to biomass. By identifying kinetic parameters of a variety of root-derived substrates, our data allow a more accurate calibration of models that consider the temporal and spatial extend of microbial growth and the mineralisation of rhizodeposits in the rhizosphere.

## Conflict of interest

None declared.

## Acknowledgements

We thank Irene Brockhaus for her technical support. This project was carried out in the framework of the priority programme 2089 “Rhizosphere spatiotemporal organization – a key to rhizosphere functions” funded by the DFG (project numbers 403635931).

## Notes

### Competing Interest Statement

The authors have declared no competing interest.

