## Supplementary Material for "Microbial utilisation of maize rhizodeposits applied to an agricultural soil at a range of concentrations"

*Running title: Microbial utilization of maize rhizodeposits*

Daniela Niedeggen[1,2], Lioba Rüger[1,2], Eva Oburger[3], Michael Santangeli[3], Mutez Ahmed[4], Doris Vetterlein[5], Sergey Blagodatsky[1,2], Michael Bonkowski[1,2]

[1]*University of Cologne, Institute for Zoology, Terrestrial Ecology, Cologne, Germany*

[2] *Cluster of Excellence on Plant Sciences (CEPLAS), University of Cologne, Germany*

[3] *University of Natural Resources and Life Sciences, Institute of Soil Research, Tulln an der Donau, Austria*

[4]*Technical University Munich –TUM, School of Life Sciences, Freising, Germany*

[5]*Helmholtz Centre for Environmental Research GmbH –UFZ, Dept. Soil System Science, Halle/Saale, Germany*

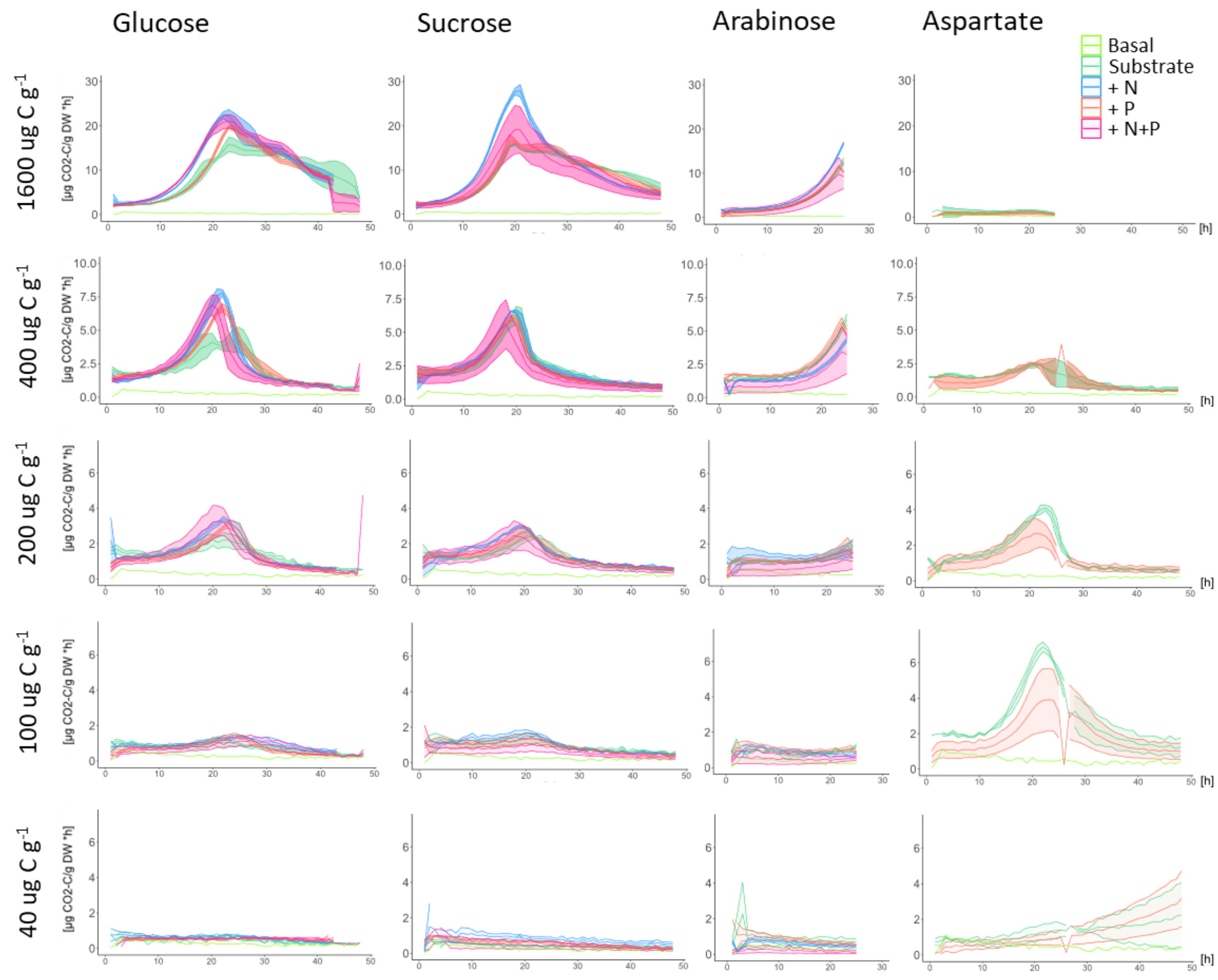


Suppl.-Fig. 1 Microbial growth in response to the sugar substrates glucose. sucrose and arabinose with addition of N, P and both; and the amino acid aspartate with addition of phosphate. The graphs are divided by concentration. Each graph shows microbial activity curves with and without N/P addition. Each curve represents the mean of 3 replicates +/- standard error.

Suppl.-Tab. 1 C-threshold and the soil Cmic specific C-threshold for microbial growth in response to the different substrates without and with nutrient supplementation.

| Substrate | | C-threshold [µg C $g^{-1}$] | Cmic specific C-threshold |
|---|---|---|---|
| **Glucose** | | 100 | 0.63 |
| | P | 40 | 0.25 |
| | N | 40 | 0.25 |
| | NP | 40 | 0.25 |
| **Sucrose** | | 100 | 0.63 |
| | P | 100 | 0.63 |
| | N | 40 | 0.25 |
| | NP | 100 | 0.63 |
| **Arabinose** | | 40 | 0.25 |
| | P | 100 | 0.63 |
| | N | 100 | 0.63 |
| | NP | 100 | 0.63 |
| **CiAcid** | | 40 | 0.63 |
| **OxAcid** | | 100 | 0.63 |
| **Aspartate** | | 40-400 | 0.25 |
| | P | 40-400 | 0.25 |
| **early Exudates** | | 400 | 2.53 |
| **late Exudates** | | 100 | 0.63 |
| **Mucilage** | | 1000 | 6.33 |

Suppl.-Tab. 2 Michaelis-Menten kinetic values ($K_m$, $V_{max}$ and $T_t$) without and with nutrient supplementation. SE represents the standard deviation of the parameter estimated by the model fit, and the p-value assesses the statistical significance of the parameter estimate. As Tt is calculated by Km and Vmax, it was only calculated if the p-values for both Km and Vmax were less than 0.1 in the model fit.

| **Substrate** | | **$K_m$** | | | **$V_{max}$** | | | **Turnovertime** |
|---|---|---|---|---|---|---|---|---|
| | | [µg C $g^{-1}$] | SE | p-value | [µg $CO_2$-C $g^{-1}$ $h^{-1}$] | SE | p-value | [h] |
| **Glucose** | | 83.22 | 34.32 | 0.032 | 2.38 | 0.25 | 0.000 | 3.16 |
| | P | 206.06 | 24.56 | 0.000 | 2.37 | 0.10 | 0.000 | 3.23 |
| | N | 281.20 | 104.40 | 0.018 | 2.66 | 0.37 | 0.000 | 2.91 |
| | NP | 875.00 | 582.30 | 0.159 | 2.74 | 0.88 | 0.009 | |
| **Sucrose** | | 161.05 | 34.38 | 0.000 | 2.53 | 0.17 | 0.000 | 3.01 |
| | P | 127.16 | 31.17 | 0.001 | 2.50 | 0.19 | 0.000 | 3.03 |
| | N | 127.79 | 57.85 | 0.046 | 2.35 | 0.32 | 0.000 | 3.22 |
| | NP | 176.80 | 121.00 | 0.168 | 2.44 | 0.55 | 0.001 | |
| **Arabinose** | | 146.54 | 52.93 | 0.017 | 1.81 | 0.20 | 0.000 | 4.20 |
| | P | 32.67 | 15.05 | 0.051 | 1.47 | 0.12 | 0.000 | 5.08 |
| | N | 75.00 | 26.82 | 0.016 | 1.77 | 0.16 | 0.000 | 4.26 |
| | NP | 232.50 | 290.70 | 0.447 | 1.36 | 0.60 | 0.055 | |
| **CiAcid** | | 19.88 | 7.85 | 0.877 | 0.87 | 0.09 | 0.000 | |
| **OxAcid** | | 7.87 | 10.83 | 0.621 | 0.42 | 0.10 | 0.000 | |
| **Aspartate** | | 11.07 | 14.85 | 0.469 | 1.54 | 0.20 | 0.000 | |
| | P | 138.10 | 30.22 | 0.655 | 0.98 | 0.24 | 0.001 | |
| **early Exudates** | | 115.50 | 47.51 | 0.035 | 2.54 | 0.29 | 0.000 | 2.98 |
| **late Exudates** | | 58.73 | 23.85 | 0.034 | 1.98 | 0.19 | 0.000 | 3.79 |
| **Mucilage** | | 3785.00 | 3524.00 | 0.296 | 10.09 | 7.17 | 0.174 | |

Suppl.-Tab. 3 Kinetic parametes $\mu_{max}$, lag-period and active fraction for soil microorganisms responding to the different substrates without and with nutrient supplementation.

| **Substrate** | | | **$\mu_{max}$** | | **Lag-period** | **active fraction** | |
|---|---|---|---|---|---|---|---|
| | + | [µg C g$^{-1}$] | | SD | [h] | [%] | SD |
| **Glucose** | | 1600 | 0,21 | 0,006 | 3,21 | 0,84 | 0,140 |
| | P | 1600 | 0,21 | 0,002 | 3,15 | 0,91 | 0,078 |
| | N | 1600 | 0,23 | 0,015 | 3,00 | 1,14 | 0,381 |
| | NP | 1600 | 0,21 | 0,003 | 2,25 | 2,16 | 0,153 |
| **Sucrose** | | 1600 | 0,26 | 0,004 | 3,84 | 0,56 | 0,079 |
| | P | 1600 | 0,25 | 0,003 | 3,54 | 0,72 | 0,069 |
| | N | 1600 | 0,23 | 0,004 | 2,14 | 2,69 | 0,436 |
| | NP | 1600 | 0,25 | 0,004 | 2,91 | 1,36 | 0,216 |
| **Arabinose** | | 1600 | 0,20 | 0,004 | 3,46 | 0,62 | 0,089 |
| | P | 1600 | 0,19 | 0,007 | 2,84 | 1,11 | 0,237 |
| | N | 1600 | 0,20 | 0,004 | 3,23 | 0,78 | 0,085 |
| | NP | 1600 | 0,19 | 0,003 | 1,70 | 3,50 | 0,343 |
| **CiAcid** | | 400 | | | | | |
| **OxAcid** | | 400 | | | | | |
| **Aspartate** | | 100 | 0,27 | 0,015 | 5,07 | 0,17 | 0,053 |
| | P | 100 | 0,25 | 0,009 | 4,38 | 0,31 | 0,240 |
| **early Exudates** | | 1600 | | | | | |
| **late Exudates** | | 1600 | | | | | |
| **Mucilage** | | 1600 | 0,18 | 0,017 | 0,96 | 6,89 | 2,633 |

SD: Standard deviation of the difference

Suppl.-Tab. 4 Abbreviations

| | |
|---|---|
| BBCH | BBCH-scale is used to identify the phenological development stages of plants |
| C | Carbon |
| Cmic | Microbial biomass |
| Km | Michaelis-Menten constant |
| KSIR | Kinetic substrate-induced respiration |
| MIRR | Maximum initial respiratory response |
| N | Nitrogen |
| P | Phosphorus |
| $T_{lag}$ | Lag-period |
| $T_t$ | Turnover time of the applied substrate by the microbial community |
| $V_{max}$ | Maximum respiration rate |
| $qCO_2$ | Specific respiration |
| $\mu_{max}$ | Maximum specific growth rate |
